# The advantages of radial trajectories for vessel-selective dynamic angiography with arterial spin labeling

**DOI:** 10.1101/683599

**Authors:** Eleanor S. K. Berry, Peter Jezzard, Thomas W. Okell

## Abstract

**Object:** To demonstrate the advantages of radial k-space trajectories over conventional Cartesian approaches for accelerating the acquisition of vessel-selective arterial spin labeling (ASL) dynamic angiograms, which are conventionally time-consuming to acquire.

**Materials and Methods:** Vessel-encoded pseudocontinuous ASL was combined with time-resolved balanced steady-state free precession (bSSFP) and spoiled gradient echo (SPGR) readouts to obtain dynamic vessel-selective angiograms arising from the four main brain-feeding arteries. Dynamic 2D protocols with acquisition times of one minute or less were achieved through radial undersampling or a Cartesian parallel imaging approach. For whole-brain dynamic 3D imaging, magnetic field inhomogeneity and the high acceleration factors required rule out the use of bSSFP and Cartesian trajectories, so the feasibility of acquiring 3D radial SPGR angiograms was tested.

**Results:** The improved SNR efficiency of bSSFP over SPGR was confirmed for 2D dynamic imaging. Radial trajectories had considerable advantages over a Cartesian approach, including a factor of two improvement in the measured SNR (p<0.00001, N=6), improved distal vessel delineation and the lack of a need for calibration data. The 3D radial approach produced good quality angiograms with negligible artifacts despite the high acceleration factor (R=13).

**Conclusion:** Radial trajectories outperform conventional Cartesian techniques for accelerated vessel-selective ASL dynamic angiography.

## Introduction

Time-resolved vessel-selective cerebral angiography provides crucial information on arterial morphology, hemodynamics and flow patterns. However, the current clinical gold standard, x-ray digital subtraction angiography, requires the use of an invasive procedure and injection of a contrast agent, resulting in some risks to the patient [1]. Non-invasive alternatives, based on Arterial Spin Labeling (ASL) MRI, show promise, but may be prohibitively slow for many clinical settings because additional measurements are required to obtain vessel-selective information from multiple arteries [2–6]. The investigation of accelerated vessel-selective ASL methods is therefore warranted, since these could allow phenomena such as occlusions, stenoses, collateral flow and blood supply to lesions to be visualized non-invasively within a clinically acceptable time frame.

Acceleration inevitably leads to loss of signal-to-noise ratio (SNR), so in order to achieve significant scan time reductions it is important that the SNR-efficiency is as high as possible. Vessel-encoded pseudocontinuous arterial spin labeling (VEPCASL) [7] is considerably more SNR-efficient than single-artery selective methods [8, 9] when multiple arteries are of interest, since all arteries contribute signal to all measurements, such that the SNR-efficiency is comparable to non-selective ASL [10]. In addition, it has been shown that VEPCASL can be combined with a balanced steady-state free precession (bSSFP) readout [11], in which transverse magnetization is “recycled” from one excitation to the next [12], unlike spoiled gradient echo (SPGR) techniques, further improving SNR-efficiency in angiographic acquisitions. However, bSSFP suffers from sensitivity to magnetic field inhomogeneity [12]. This can lead to significant ASL signal loss when imaging is performed over large regions where B_0_shimming is more challenging [11].

If high SNR-efficiency can be achieved, then acceleration through undersampling should be possible whilst maintaining reasonable image quality. However, undersampling with traditional Cartesian trajectories results in strong, coherent ghosting artifacts unless the data are reconstructed with parallel imaging algorithms [13, 14]. This requires additional scan time for calibration and results in noise amplification, which is typically most severe at the center of the head where many cerebral vessels of interest reside. Radial acquisition schemes can be angularly undersampled, resulting in aliasing artifacts that take the form of noise-like signal variations and streaks [15]. Undersampled radial readouts are particularly applicable for cerebral angiography because the signal is sparse, so streak artifacts are minimal and often relegated to the image periphery [16, 17]. Radial trajectories also result in more manageable flow or respiration artifacts than Cartesian acquisitions [15], can give more accurate timing information [18], and have previously been used with success for 3D non-vessel-selective ASL angiography [18–23].

In this work we compare undersampled radial and Cartesian acquisition schemes for the acquisition of accelerated dynamic vessel-selective ASL cerebral angiograms. In the context of rapid 2D dynamic protocols we also compare bSSFP and SPGR, as bSSFP artifacts can be scan-time and trajectory dependent [24]. We then explore the potential for 3D whole brain acquisitions: over a large imaging region bSSFP artifacts are problematic and the required acceleration factors are so high that conventional Cartesian approaches are not feasible, so we instead demonstrate the potential of a SPGR radial approach. This work follows on from a previously presented conference abstract [25].

## Materials and Methods

### Pulse Sequence Design and Image Reconstruction

The VEPCASL dynamic angiographic pulse sequence is shown schematically in Figure 1. Both undersampled Cartesian and radial trajectories were implemented. For the Cartesian case, uniform undersampling was performed to allow a conventional parallel imaging reconstruction. The 2D radial trajectory used in this work consisted of full radial spokes with uniform azimuthal spacing [15] (see Fig. 1b). The 3D radial trajectory also used full radial spokes with starting points evenly distributed across a hemisphere [26] (Fig. 1c), similar to previous studies [19, 27]. In all cases, multiple adjacent k-space lines (a “segment”) were acquired repeatedly following each VEPCASL preparation, as per previous implementations [11]. The order of segment acquisition here was sequential, i.e. the whole of k-space was progressively acquired for a given VEPCASL cycle before moving on to the next cycle [11]. Temporal information was gathered by acquiring a given segment multiple times following labeling, to track the flow of labeled blood through the vasculature. The acquisition could be triggered through cardiac gating via a pulse oximeter. For bSSFP protocols, a series of 20 RF pulses with linearly increasing flip angles were used prior to imaging to minimize transient signal oscillations during the approach to steady state, as used previously [11].

**Fig 1:**
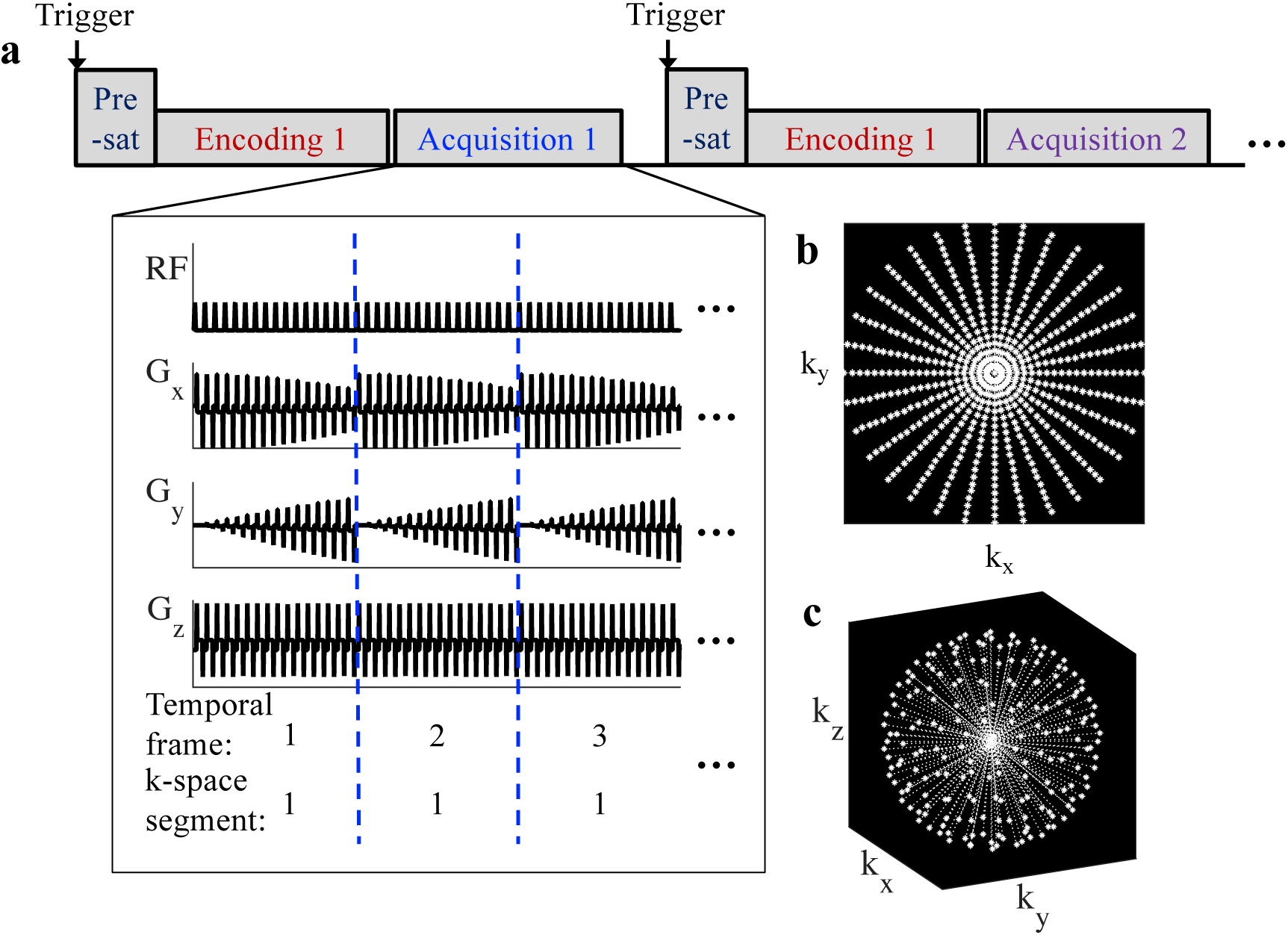
a) Sequence schematic for VEPCASL dynamic angiography with sequential segment ordering. Acquisition number corresponds to the number of the k-space segment acquired. Encoding number refers to the applied (VE)PCASL labeling cycle. The start of the acquisitions can be triggered using a pulse oximeter, which is followed by a pre-saturation (“pre-sat”) module. For the bSSFP readout, a series of 20 RF pulses with linearly increasing flip angles was used to stabilize the signal before acquisition began. Example gradient waveforms for the 2D radial trajectory are also shown. b) Example k-space trajectory for a 2D radial acquisition. c) Example k-space trajectory for a 3D radial acquisition.

Vendor image reconstruction based on gridding [15] or GRAPPA [14] was used for the 2D radial and Cartesian acquisitions, respectively. 3D radial data were reconstructed offline in Matlab (Mathworks, Natick, Massachusetts, USA) using the adjoint operator to the non-uniform fast Fourier transform [28, 29], which is akin to regridding, with sampling density compensation calculated as per [30]. In all cases separate images from each coil were reconstructed before combining using a sum of squares technique [31].

### Experiments

#### Scan Protocols

All scans were performed on a 3T TIM Verio system (Siemens Healthineers, Erlangen, Germany), with a 32-channel head coil. All subjects were scanned under a technical development protocol approved by local ethics and institutional committees.

Pre-scans included three-dimensional multislab time-of-flight (TOF) angiography to allow selection of a labeling plane and localization of the four main brain-feeding arteries in the neck (voxel size = 0.8 × 0.8 × 1.3 mm). A field map of the labeling plane (voxel size = 0.9 × 0.9 × 2.0 mm, TR = 200 ms, TE1 = 5.19 ms, TE2 = 6.19 ms, acquisition time = 1 minute) was acquired and a set of eight encodings, similar to Okell et al. [10], calculated using the optimized encoding scheme [32], including a correction for any off-resonance effects within the labeling plane [33].

These encodings were used to perform vessel-selective angiography of the right and left internal carotid and vertebral arteries using a unipolar VEPCASL [34] pulse train of duration 800 ms with other labeling parameters as per Berry et al. [32]. Cardiac gating has been shown to improve bSSFP imaging to a certain degree [35] so was employed during all bSSFP scans. An iterative shimming procedure (automatic shimming performed three times) was used to minimize B_0_ inhomogeneity within the imaging region for the bSSFP scans. For dynamic 2D imaging, the in-plane resolution was fixed at 0.9 × 0.9 mm with a slice thickness of 50 mm to encompass the circle of Willis and nearby arterial branches. A temporal resolution of approximately 100 ms was chosen here, in line with Yan et al. [36], to provide good temporal fidelity whilst constraining the total scan time. For dynamic 3D imaging, non-selective RF pulses were used to excite the whole head. The spatial resolution was set to 1.17 mm isotropic and the temporal resolution to 191 ms to keep the total scan time within reasonable limits. Other imaging parameters varied between experiments, as detailed below.

#### Comparison of bSSFP and SPGR for 2D Dynamic Angiography

Preliminary sequence testing was performed to confirm the expected benefits of bSSFP over SPGR readouts when combined with rapid 2D Cartesian and radial trajectories. An accelerated Cartesian acquisition was compared to a time-matched radially undersampled protocol. A further undersampled radial scheme that can achieve acquisition times of less than one minute was also performed to demonstrate the potential for further acceleration with a radial trajectory. Cartesian scan times included the time required for the acquisition of GRAPPA calibration data.

For this comparison, a healthy male subject (31 years) was scanned. Six different readout strategies, three SPGR and three bSSFP, were used, with acquisition parameters listed in Table 1 (protocols 1-6). The bSSFP acquisitions were cardiac gated, so the acquisition times listed are predicted times based on a 1 second cardiac period.

**Table 1:** Readout parameters for each VEPCASL angiography protocol ^*a*^ *Cardiac cycle dependent*

| Readout parameters | 2D |  |  |  |  |  | 3D |
| --- | --- | --- | --- | --- | --- | --- | --- |
|  | *SPGR* |  |  | *bSSFP* |  |  | *SPGR* |
| *Protocol number* | *1* | *2* | *3* | *4* | *5* | *6* | *7* |
| Trajectory | Cartesian | Radial | Radial | Cartesian | Radial | Radial | Radial |
| # k space lines | 56 | 84 | 56 | 56 | 80 | 60 | 2048 |
| Acceleration factor | 4 | 4.2 | 6.3 | 4 | 4.4 | 5.9 | 12.6 |
| Approximate<sup>a</sup> total acquisition time (s) | 88 | 90 | 64 | 63 | 63 | 50 | 960 |
| Readout flip angle (°) | 8 | 8 | 8 | 40 | 40 | 40 | 8 |
| # frames | 7 | 8 | 8 | 6 | 6 | 6 | 5 |
| Temporal resolution (ms) | 101.92 | 93.94 | 93.94 | 101.8 | 101.8 | 101.8 | 191.4 |
| k-space lines per ASL preparation per frame | 14 | 14 | 14 | 20 | 20 | 20 | 32 |
| TR (ms) | 7.28 | 6.71 | 6.71 | 5.09 | 5.09 | 5.09 | 5.98 |
| TE (ms) | 3.59 | 3.58 | 3.58 | 2.55 | 2.55 | 2.55 | 2.67 |
| Readout bandwidth (Hz/Px) | 302 | 302 | 302 | 475 | 475 | 475 | 300 |
| Matrix size | 224 x 224 |  |  |  |  |  | 128<sup>3</sup> |
| Voxel size (mm) | 0.9 x 0.9 x 50 |  |  |  |  |  | 1.17<sup>3</sup> |
| Image FOV (mm) | 200 x 200 x 50 |  |  |  |  |  | 150<sup>3</sup> |

The SPGR and bSSFP sequence parameters were chosen to ensure, firstly, similar undersampling factors and temporal resolution between the two readout types and, secondly, to optimize SPGR/bSSFP image quality. The choice of flip angle for SPGR sequences is a tradeoff between high initial signal and the level of signal attenuation that is acceptable for the duration of the scan. By the end of the imaging readout period the SPGR parameters chosen here, with relatively short TR to keep the imaging times short, result in signal attenuation due to the imaging pulses of approximately 66%, calculated as per Okell et al. [37]. Note that the minimum TEs for SPGR Cartesian and radial acquisitions were different, as no phase encoding is required with a radial trajectory. This resulted in small differences in TR and temporal resolution. To minimize banding artifacts in the bSSFP images the TR was kept short and the flip angle chosen to be in line with optimization of previous bSSFP VEPCASL acquisitions [11].

#### Comparison of Accelerated Cartesian and Radial 2D Trajectories

Six healthy male subjects (mean age 30.3 ± 4.5 years) were scanned to compare accelerated Cartesian and radial trajectories for dynamic 2D angiography. As a result of the SPGR and bSSFP experiments detailed above, the scan protocol included only the three different bSSFP acquisition strategies: accelerated Cartesian, time-matched radial and further undersampled radial trajectories (Table 1, protocols 4-6).

#### Dynamic 3D Acquisition

Extending the VEPCASL angiography acquisition scheme to a dynamic 3D readout with whole head coverage requires an acceleration factor too large to be supported by conventional Cartesian trajectories with parallel imaging. However, such an acquisition is feasible with an undersampled radial trajectory. To demonstrate the potential of this approach, one healthy volunteer (female, age 34 years) was scanned with VEPCASL labeling parameters as per the dynamic 2D protocols and readout parameters as shown in Table 1 (protocol 7). Spatial and temporal resolution were compromised (1.17 mm isotropic and 191 ms, respectively) relative to the 2D protocols to achieve a reasonable scan time of 16 minutes, corresponding to an undersampling factor of 12.6.

### Image Analysis

All VEPCASL images were analyzed using a Bayesian maximum *a posteriori* (MAP) method [38] to separate out vessel-specific information. In order to quantitatively compare the 2D accelerated Cartesian and radial approaches the SNR in the vessel-selective angiograms for each subject was calculated. Vessel-specific signal masks were generated as follows:

1. Signal from outside the brain and the scalp was removed using a manually drawn mask.
2. A signal threshold was applied to voxels within the brain mask across all time points. Only voxels with signal above the 80^th^ percentile of the signal across all acquisitions were included. This was empirically found to reduce the inclusion of voxels outside the main brain-feeding arteries and ensure that the same voxels were compared across acquisitions.
3. Vessel-specific signal masks were created according to the dominant feeding artery of each voxel (i.e. the artery contributing the highest signal) meeting the signal threshold. If an artery was not the dominant feeding artery to any voxels, it was excluded from the comparison.

A noise region-of-interest (ROI), of around 40 voxels, was selected from the center of the brain of each subject, avoiding any arteries, to assess noise close to vessels of interest. The mean signal was calculated within the vessel mask for each artery and the standard deviation of the noise calculated from the noise ROI. The ratio yielded the SNR, which was compared across the different protocols using paired t-tests.

This approach results in the measured “noise” containing contributions from both true noise and aliased signal due to radial undersampling or imperfect parallel imaging reconstruction. However, the choice of a noise ROI close to the arteries of interest inside the brain ensures that the measured noise represents the signal fluctuations above which the ASL signal must rise in order to be clearly visible.

## Results

### Comparison of bSSFP and SPGR for 2D Dynamic Angiography

Across subjects, scan times of approximately one minute and below were achieved for the dynamic 2D protocols, dependent on the pulse rate of the individual subject. Figure 2 demonstrates the differences in image quality obtained between SPGR and bSSFP across the different trajectory types in the subject used for initial 2D sequence testing. The subtraction images of the non-selective tag and control encoding cycles show that the bSSFP data were visibly less noisy than their SPGR counterparts. The average measured SNR of these bSSFP images relative to the equivalent SPGR acquisitions was 2.0. The posterior circulation in this subject contained a lower blood signal than the anterior circulation, making it difficult to see above the noise in the SPGR data, but much clearer visualization is achieved with the bSSFP approach. It is also worth noting the visible noise amplification and residual signal aliasing, which can be seen within the brain for the accelerated Cartesian protocol compared with the time-matched radial protocol. Radial streaking artifacts become more pronounced at higher acceleration factors, although these are more evident outside the brain. No obvious differences in flow-related artifacts were observed between Cartesian and radial acquisitions. As a result of these observations, a bSSFP acquisition was used for the remainder of the dynamic 2D experiments.

**Fig 2:**
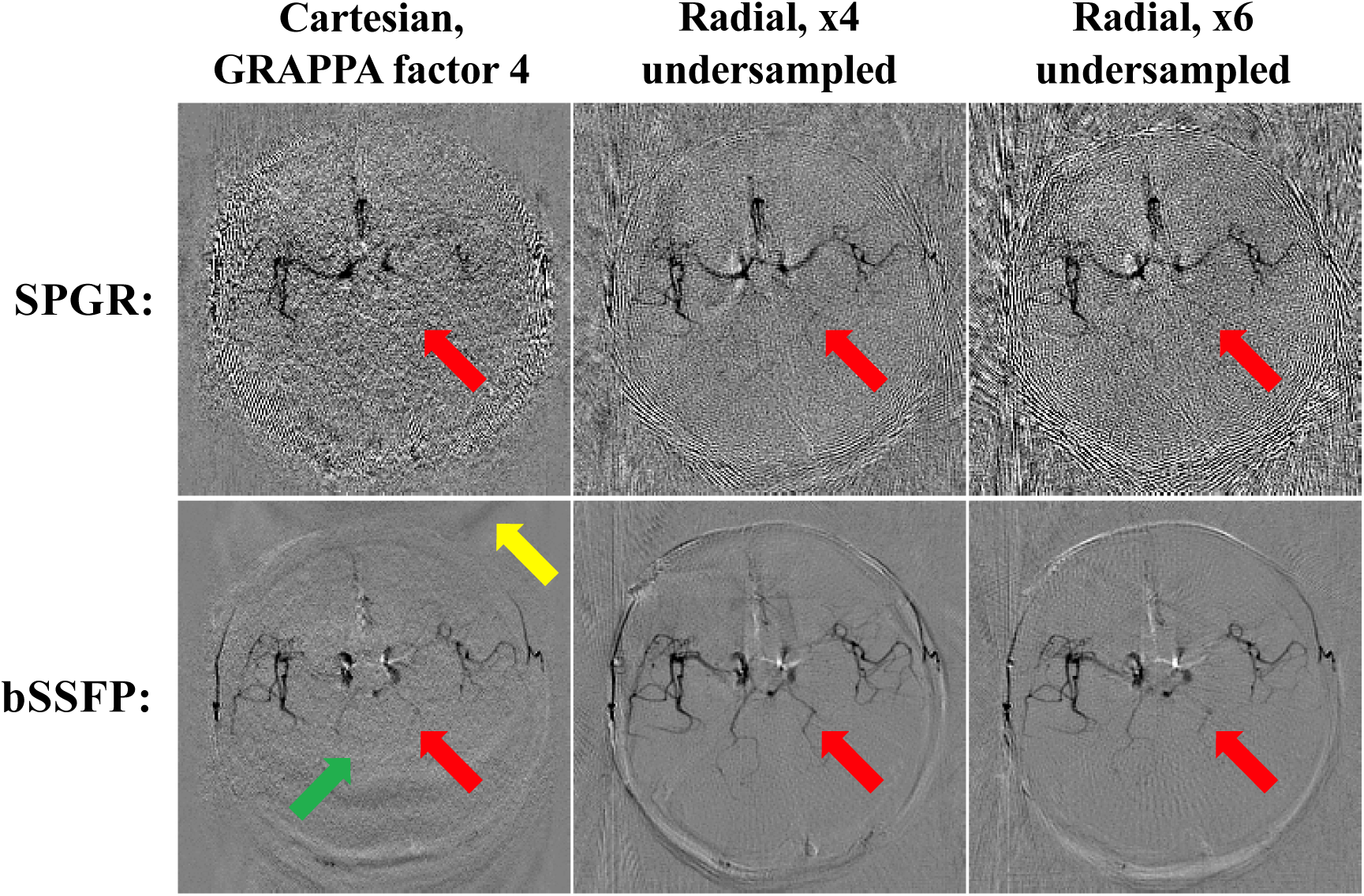
Comparison of SPGR and bSSFP: Subtraction of the non-selective tag and control encoding cycles (giving dark blood) for the first time frame of the SPGR and bSSFP data sets across all three trajectories. Note the visibly improved SNR obtained with bSSFP, allowing clearer delineation of fine vessels, such as those in the posterior circulation (red arrows). Some spatially varying noise amplification (green arrow) and residual aliasing (yellow arrow) are also visible in the accelerated Cartesian images.

### Comparison of Accelerated Cartesian and Radial 2D Trajectories

Figure 3 shows the flow of blood through the vessels of one of the subjects for each of the different 2D k-space trajectories. Note that due to the use of a relatively long VEPCASL pulse train, the first frame shows most of the vasculature filled with labeled blood. For all acquisitions the arteries are relatively well visualized and movement of the blood from the central circle of Willis to the peripheral circulation over time can be seen. The posterior circulation has a delayed arrival time, which is consistent with previous angiographic [37] and perfusion [39] data. A greater degree of noise is apparent in the center of the Cartesian images compared to the time-matched radial data. As expected, increasing the radial undersampling leads to lower apparent SNR, although most of the vessels are still clearly visualized.

**Fig 3:**
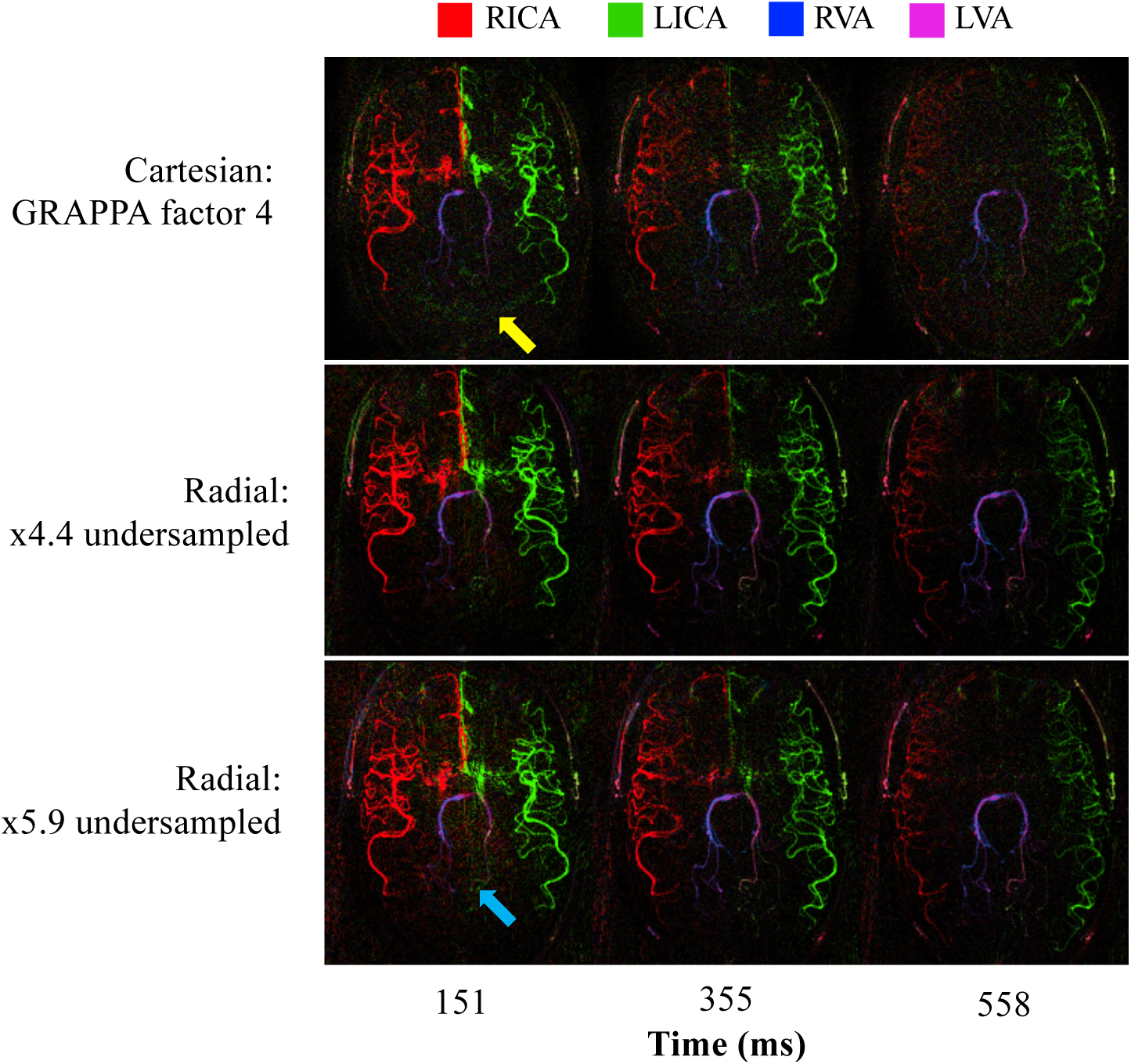
Selected frames from dynamic 2D vessel-selective angiograms using bSSFP for a single subject across the three acquisition schemes. Comparable dynamic information is obtained in all cases. However, the accelerated Cartesian data is noticeably noisier and contains some residual aliased signal in some regions (yellow arrow). Noise-like aliasing in the radial data becomes more noticeable at a higher undersampling factor (blue arrow). The arterial source of the blood signal is shown in color (see legend). The displayed time is that after the end of the VEPCASL pulse train.

Figure 4a shows the time-averaged vessel-selective images for a single subject across the three 2D acquisition schemes. The superior SNR of the radial images is apparent in the vessel definition and visualization of narrow vessel branches. Minor streak artifacts are in evidence at the edges of the radially acquired images, particularly for the more heavily undersampled scan. There are also artifacts at the center of all the images, including disruption of the proximal middle cerebral artery signal, which could be due to pulsatility or residual B_0_ inhomogeneity in this area (yellow arrows).

**Fig 4:**
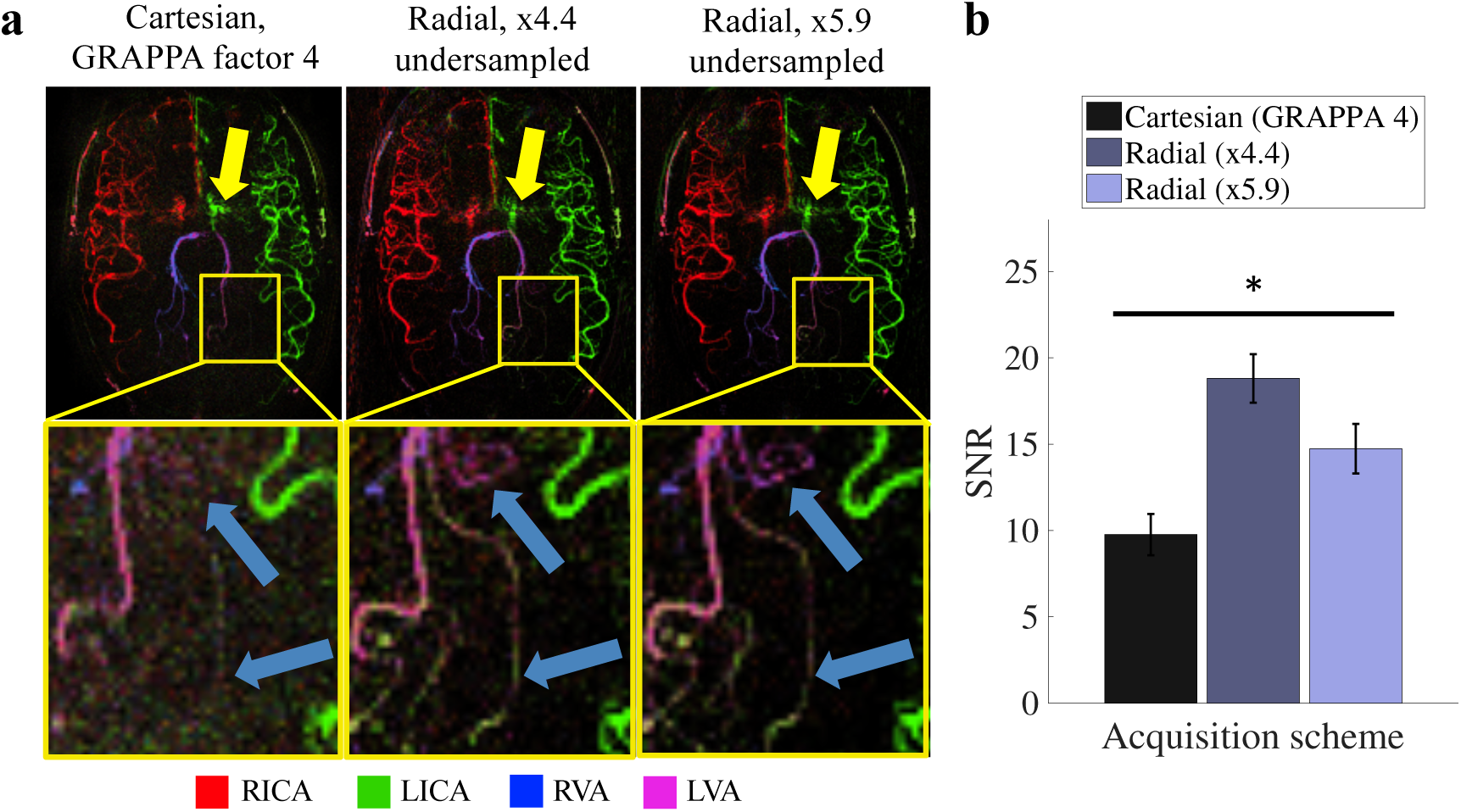
2D image quality comparison: a) Time averaged vessel-selective images for a single subject (top row). There is some artifact in the proximal arteries, perhaps due to pulsatility or B_0_ inhomogeneity (yellow arrows). An enlarged section of the images (bottom row) demonstrates that the noise amplification from GRAPPA obscures fine vessel details that are visible in the radial acquisitions (blue arrows); b) Mean SNR across vessels for all acquisition schemes. All acquisitions have significantly different SNR to each other (\**p* < 10^-5^). Error bars represent the standard error.

The SNR for all acquisitions is shown in Figure 4b, demonstrating that the accelerated Cartesian protocol resulted in approximately half the SNR of the time-matched radial acquisition. Increasing the radial undersampling from x4.4 to x5.9 resulted in a modest SNR decrease of approximately 20%, but this still out-performed the slower Cartesian acquisition. According to paired t-tests the mean SNR of each acquisition strategy is significantly different to the others (p < 10^-5^).

### Dynamic 3D Acquisition

Temporal average maximum intensity projections (MIPs) of the dynamic 3D radial SPGR VEPCASL angiogram are shown in Figure 5. The SNR benefit of a 3D acquisition compensated for the lower SNR efficiency of the SPGR sequence used in this case, resulting in good vessel definition with minimal noise across the whole brain. Signal aliasing and artifacts resulting from the high undersampling factor (x13) did not appear to impede image quality. In addition, the visualization of the proximal arteries, including the MCAs, was not hampered by pulsatility or B_0_ inhomogeneity, as was apparent in the 2D bSSFP data.

**Fig 5:**
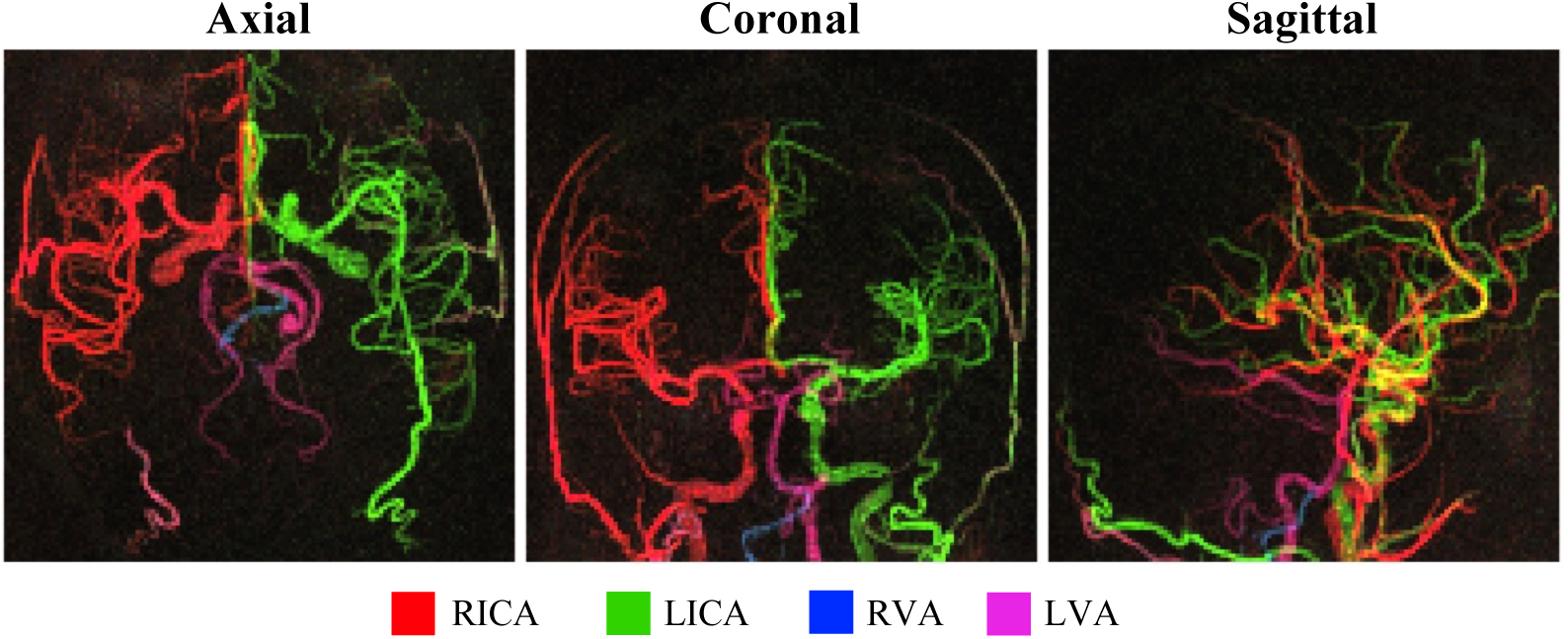
Dynamic 3D VEPCASL SPGR angiogram, shown as axial, coronal and sagittal MIPs of the temporal mean image. Note that the yellow color in the sagittal view is a result of the image view point, where arteries containing blood from the RICA and LICA overlap, rather than mixed blood supply. Note the lack of proximal artery artifacts, good vessel delineation and minimal background signal.

Axial MIPs of the individual time frames are shown in Figure 6, demonstrating the dynamic information available with this acquisition scheme. As with the dynamic 2D data, delayed washout of the bolus is apparent in the posterior circulation and sufficient signal is retained in later time frames to visualize the more distal vessels.

**Fig 6:**
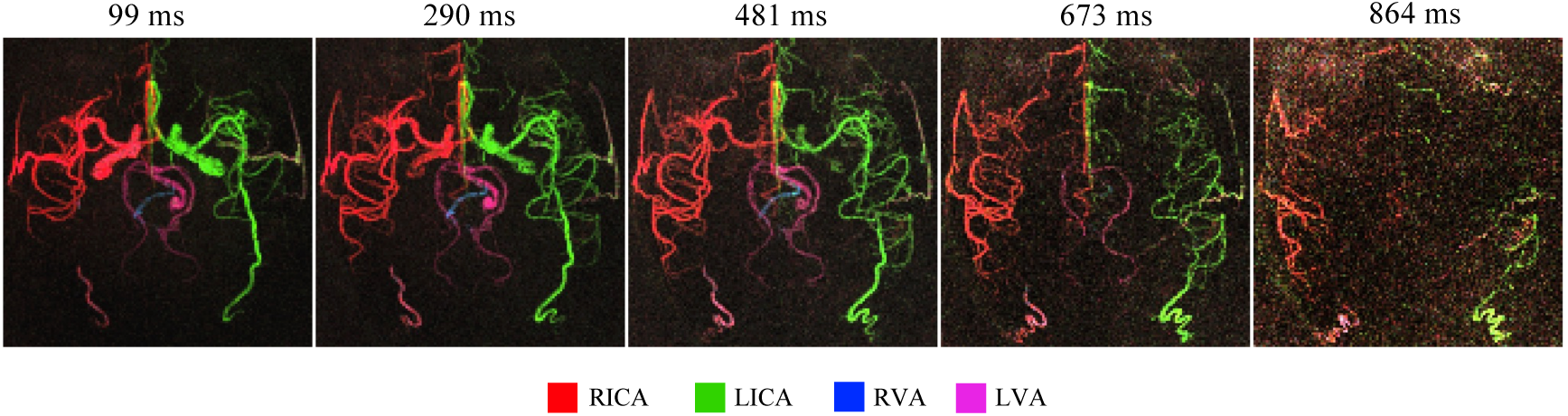
Axial MIP of each frame of the dynamic 3D VEPCASL SPGR angiogram shown in Figure 5, demonstrating the temporal dynamics of the signal. The time above each frame is given as the delay after the end of the VEPCASL pulse train.

## Discussion

The ability to accelerate image acquisition is crucial for vessel-selective dynamic angiography with ASL where additional measurements are required beyond non-selective approaches. In this study we have shown that radial trajectories have considerable benefits over conventional Cartesian techniques in this context. These include: a) the lack of need to acquire calibration data for parallel imaging, allowing all the available scan time to be used for imaging; b) higher apparent SNR and distal vessel visibility; and c) no noise amplification in the middle of the brain or residual aliasing arising from the parallel imaging reconstructions. The sparse nature of the angiograms meant that radial undersampling artifacts were relatively benign and most prominent at the edges of the field of view, away from the vessels of interest. The ability of radial trajectories to accelerate the acquisition was demonstrated by the acquisition of 2D vessel-selective angiograms in less than one minute. In 3D, even greater acceleration factors are possible, allowing the first demonstration of whole-brain vessel-selective dynamic angiography with a 3D radial trajectory within a reasonable scan time (16 minutes).

For vessel-selective dynamic 2D angiography, it was found that bSSFP resulted in images with higher SNR than SPGR images, as expected from previous work [11]. No increase in flow-related artifacts were observed using radial trajectories compared to a conventional Cartesian approach. The Cartesian readout with high GRAPPA factor suffered from reduced SNR towards the center of the brain, around vessels of interest, which is to be expected due to the higher g-factor in this region [40]. Radial undersampling, however, led to increased noise or signal aliasing which was most apparent at the edges of the FOV, away from these vessels.

As a result, the time-matched radial acquisition had a two-fold better SNR than the Cartesian acquisition within the major cerebral vessels. Increasing the undersampling factor for the radial acquisition to achieve scans times of less than one minute resulted in vessel-selective angiograms with increased streak artifacts, but the SNR was still higher and the distal arteries still better delineated than in accelerated Cartesian acquisitions (Figure 4). Image quality could perhaps be further improved through the use of parallel imaging in the radial image reconstruction [41]. As well as increased noise amplification in the centre of the brain, accelerated Cartesian acquisitions also require additional calibration data, the measurement of which can take up a significant proportion of the scan time in this kind of rapid acquisition, further reducing the SNR efficiency.

For a 3D acquisition, it was not possible to use bSSFP due to the greater B_0_ inhomogeneity present across the whole brain, which has previously been shown to be problematic for ASL angiography [11]. However, the increased SNR of a 3D acquisition still allowed good image quality to be obtained using an SPGR readout, despite the high undersampling factor required (x13). This degree of undersampling was not feasible using a conventional Cartesian parallel imaging reconstruction, although in future work the application of more advanced approaches such as blipped-CAIPI sampling [42] may allow higher acceleration factors. The use of a true 3D radial approach in this study allowed the benefits of radial trajectories to apply in all three dimensions and gave whole-brain coverage within a reasonable scan time. However, particularly when only a reduced field-of-view is of interest, comparison of this approach with a stack-of-stars trajectory [21, 23] would be useful in future work, as well as validation in a greater number of subjects.

The results of this study are consistent with previous work that has used various radial approaches for non-selective ASL angiography. For example, Koktzoglou et al. [19, 22] and Wu et al. [20] have demonstrated the excellent image quality achievable with radial trajectories, although no direct comparison with a matched Cartesian approach was performed. Song et al. [21] and Cong et al. [23] showed comparable image quality with an accelerated stack-of-stars trajectory combined with a k-space filtering approach compared to a fully sampled Cartesian acquisition. Wu et al. [18] also showed the improved timing information obtained from a radial approach compared to a Cartesian method in a digital phantom, although no experimental comparison was performed. In this study we built upon this prior work by performing a direct experimental comparison of time-matched accelerated Cartesian and radial trajectories in the context of vessel-selective ASL angiography, where acceleration is particularly important, and utilize the benefits of a true 3D radial trajectory to perform whole-brain vessel-selective dynamic ASL angiography.

One downside of using bSSFP for the 2D acquisitions in this study was the appearance of artifacts in the proximal arteries in some subjects (Figure 4), which were not observed in the SPGR data (Figures 2, 5 and 6). These may be a result of pulsatile blood and/or cerebrospinal fluid flow, or the presence of B_0_ inhomogeneity. Similar artifacts were not observed in previous work on VEPCASL angiography with bSSFP [11]. One possible explanation is the difference in the VEPCASL pulse train duration: here the shorter tag duration was approximately the same as the cardiac cycle, perhaps giving poorer image quality in the initial frames due to data acquisition during systole. In addition, the TR used here was about 20% longer than the previous study and data were acquired on a different scanner, so the increased sensitivity to off-resonance effects combined with a different shimming set-up could have led to artifacts in this region. Further investigation of this phenomenon is necessary before such a protocol could be deployed clinically. However, these artifacts were not specific to the trajectory, so we anticipate that the benefits of radial imaging over conventional Cartesian methods found in this study should generalize to other protocols where such artifacts have been minimized.

The measure of SNR used in this study is somewhat qualitative. It allowed aliased signal from undersampling to be included, which affects estimates of both the vascular signal and the noise measure. The effect on the mean signal was likely to be small, as the aliasing within the brain was not severe. However, the aliased signal will contribute to the apparent noise in ROIs close to the arteries in such a way as to include information on signal fluctuations not due to local vascular signal, but which will still hinder image interpretation, and are therefore important to consider. In addition, the SNR metric was used here to allow comparison between protocols, rather than to determine the absolute image SNR.

Whilst the dynamic 2D protocols described here can take as little as one minute to acquire, in this proof-of-concept study the set-up time, including TOF and field map data of the labeling plane and use of an iterative shimming procedure, all contributed to the total scan time. In future work, scan time could be minimized by: a) fixing the location of the labeling plane relative to an anatomical landmark [43] or using a planning-free approach [44], removing the need for a TOF acquisition; b) removing the field map scan used for off-resonance correction, since magnetic field inhomogeneity is typically small at this level in the neck, even when shimming is only performed over the imaging region [32]; and c) reducing the sensitivity of bSSFP to field inhomogeneity by using a shorter TR and reduced flip angle, or by moving to a lower field strength (e.g. 1.5 T), or switching to an SPGR approach, thereby removing the need for an iterative shim. However, since these set-up procedures were matched between the acquisitions, these factors should not have affected the observed improvements in image quality obtained with a radial trajectory compared to a conventional Cartesian approach.

The next stage of this work would be to increase the robustness of the 2D bSSFP approach in proximal vessels and explore the potential for further scan time reductions in the 3D SPGR acquisition. Comparisons with a gold standard, such as x-ray-based methods, and assessment through clinician scoring would help to establish the usefulness of these approaches in a clinical setting. The inclusion of the protocol on a patient population would better evaluate its ability to provide dynamic vessel-selective information in diseased cerebral vasculature.

Additionally, the focus of this work has been time-resolved angiograms with vessel-specific information for four arteries, but it could easily be extended to more than four vessels, albeit with the requirement for a longer scan time. The optimized encoding scheme technique with off-resonance correction [33] is able to generate a minimal number of optimized encodings for any number of vessels. Consequently, the number of encoding cycles can be tailored to the number of vessels of interest and kept to a minimum. The result of this would be relatively short total scan times for many-vessel dynamic angiographic information. In addition, further acceleration could be achieved through the use of advanced image reconstruction techniques such as compressed sensing [45], which are well suited to sparse angiographic data and radial trajectories [46]. Additional flexibility in the reconstruction could also be provided through the use of golden ratio spoke ordering [47, 48], allowing the temporal resolution and undersampling factors to be chosen retrospectively.

## Conclusions

In this study we have demonstrated the benefits of radial trajectories over Cartesian methods for non-invasive vessel-selective dynamic angiograms of the main cerebral vessels. This allowed 2D dynamic angiograms to be obtained in one minute or less, and highly accelerated whole-brain 3D dynamic angiograms to be acquired in 16 minutes, whilst maintaining good image quality.

## Author Contributions

Berry: Study conception and design; Acquisition of data; Analysis and interpretation of data; Drafting of manuscript

Jezzard: Study conception and design; Critical revision

Okell: Study conception and design; Analysis and interpretation of data; Drafting of manuscript; Critical revision

## Compliance with Ethical Standards

### Funding

This work was supported by the Engineering and Physical Sciences Research Council, The Dunhill Medical Trust [grant number OSRP1/1006] and the Royal Academy of Engineering [grant number RF/132]. The Wellcome Centre for Integrative Neuroimaging is supported by core funding from the Wellcome Trust [grant number 203139/Z/16/Z] and PJ is supported by the National Institute for Health Research Oxford Biomedical Research Centre.

### Ethical approval

All procedures performed in studies involving human participants were in accordance with the ethical standards of the institutional research committee (Oxford University Clinical Trials and Research Governance office, Technical Development SOP FMRIB_004_V4) and with the 1964 Helsinki declaration and its later amendments or comparable ethical standards.

### Informed consent

Informed consent was obtained from all individual participants included in the study.

